# Bacteria: A novel source for potent mosquito feeding – deterrents

**DOI:** 10.1101/424788

**Authors:** Mayur K. Kajla, Gregory A. Barrett-Wilt, Susan M. Paskewitz

## Abstract

Antibiotic and insecticidal bioactivities of the extracellular secondary metabolites produced by entomopathogenic bacteria belonging to genus Xenorhabdus have been identified; however, their novel applications such as mosquito feeding-deterrence have not been reported. Here, we show that a mixture of compounds isolated from *Xenorhabdus budapestensis in vitro* cultures exhibits potent feeding-deterrent activity against three deadly mosquito vectors: *Aedes aegypti, Anopheles gambiae* and *Culex pipiens*. We further demonstrate that the deterrent-active fraction isolated from replicate bacterial cultures is consistently highly enriched in two modified peptides identical to the previously described fabclavines, strongly suggesting that these are molecular species responsible for feeding-deterrence. The mosquito feeding-deterrent activity in the fabclavines-rich fraction is comparable to or better than that of N, N-diethyl-3-methylbenzamide (also known as Deet) or picaridin in side-by-side assays. Our unique discovery lays the groundwork for research into biologically derived, peptide-based low molecular weight compounds isolated from bacteria for exploitation as mosquito repellents and feeding-deterrents.

## Introduction

Secondary or specialized metabolites (*1*) are a chemically diverse group of organic compounds produced by some microbes, plants and animals (*2*). Generally considered non-essential for growth and development, secondary metabolites play other roles such as conferring protection against varied environmental risks. Many secondary metabolites of microbial or plant origin have been exploited for myriad applications in the pharmaceutical industry including antibiotics, chemotherapeutic drugs, immune suppressants and other medicines (*2*). Secondary metabolites produced by *Xenorhabdus*, a group of bacteria that symbiotically associate with entomopathogenic nematodes, exhibit a range of antibiotic, antifungal and insecticidal activities (*3-9*).

Genome studies (*6, 10*) suggest that there is an enormous range of chemical diversity and bioactivity which still remains to be explored and may lead to discovery of novel bioactive compounds. In *Xenorhabdus*, genome mining has uncovered gene clusters such as polyketide synthases or non-ribosomal peptide synthases predicted to participate in synthesis of several compounds of unknown biological functions (*11*). The products encoded by these gene clusters exhibit diverse and structurally complex chemistries including the formation of modified peptides, polyketides or unique hybrids of both. One example produced by *X. budapestensis* (Xbu) is a unique class of hybrid compounds called fabclavines, that share a peptide-polyketide-polyamino backbone (*11*). Interestingly, fabclavines also exhibit a broad range of bioactivities similar to other Xenorhabdus secondary metabolites (*3, 4, 11, 12*). In addition to antibiotic and insecticidal activities, culture supernatants of *X. nematophila* and *Photorhabdus luminescens* were reported to deter feeding of ants, crickets, and wasps but neither study isolated or identified the active compound/s (*13, 14*). These studies suggested that these or related compounds might act as **m**osquito **f**eeding **d**eterrents (hereafter referred as MFDs).

Repellents (topical and area) can provide protection from mosquito (and other blood-feeding insects) bites and thus from the disease agents that they transmit while feeding (*15, 16*). Mosquitoes transmit pathogens that cause devastating human diseases, including dengue, Chikungunya, West Nile and Zika viral infections that continue to affect millions of people worldwide. Limiting the impact of mosquito-borne diseases is an important goal for global public health agencies and the use of mosquito repellents is one important tactic. Among EPA-registered topical insect repellents, a majority of the over 500 products contain the active ingredient N, N-diethyl-3-methylbenzamide (also known as Deet; www.epa.gov/insect-repellents/skin-applied-repellent-ingredients), which is the most widely used and effective repellent against mosquitoes and other disease vectors. Other commercially successful (*17*) insect repellents include IR3535 (Insect Repellent 3535, EBAAP, a derivative of β-alanine (*18*)), (p)icaridin (and other piperidine derivatives), (*19*) and para-menthane-3,8-diol (distilled from *Eucalyptus citriodora*). Historically, research into biologically-derived Deet alternatives has primarily focused on plant metabolites. Despite the exploitation of many genera for pharmaceutical exploration, bacteria have thus far remained unexplored in the search for insect repellent chemistries.

In a screen for repellent activities produced by *Xenorhabdus*, we observed that Xbu extracts deterred adult female *Aedes, Anopheles* and *Culex* mosquitoes from feeding on an artificial diet in *in vitro* feeding experiments. The isolated MFDs produced by these bacteria are composed of two low-molecular weight compounds identical to previously reported molecules called fabclavines (*11*). In this report, we present data on purification, identification and bioactivities of fabclavines as potent mosquito feeding-deterrents. Our discovery adds bacteria as a significant, potential source of novel MFDs and lays the groundwork for future exploration of bacterially derived secondary metabolites as feeding deterrents for other pest insects.

## Results and discussion

### In vitro membrane feeding system as a screening assay for determining mosquito feeding-deterrent activity in Xbu extracts

In the context of insect repellents, a deterrent is defined as “something that inhibits feeding or oviposition when present in a place where the insect would, in its absence, feed, rest or oviposit” (*20*). In this report, we define compounds produced by Xbu as MFDs because in the presence of Xbu compounds female mosquitoes failed to feed on the food source described in this section.

Among methodologies to evaluate mosquito deterrents/repellents, *in vitro* assays offer important advantages. Due to the risk associated with accidental exposure to infectious agents during standard arm-in-cage assays (*21, 22*), and because of inconsistency in results among test subjects due to varying mosquito attraction to human subjects (*23*), researchers have developed artificial feeding systems. Blood is contained in a warming chamber and accessed by adult female mosquitoes through a natural or artificial membrane. Membrane feeders provide flexible systems that can be modified in terms of feeding solutions (*24-26*), membrane types (*24, 25, 27*), and solution temperature (*24, 25, 28*). Such systems eliminate the need to expose animals (*29*) or human volunteers, an especially important consideration when the compounds in the initial screening phases are of unknown toxicological or dermatological properties. Thus, we chose to modify our existing Hemotek membrane feeding system (Discovery Workshops, Accrington, UK) to screen feeding-deterrent compounds produced by Xbu bacteria. The main features of this system (Fig. 1) included 1) easy set-up, 2) minimal laboratory space requirements, 3) a thermostat regulated temperature source eliminating the need for a water heater and circulation pump, and 4) capacity for up to five screening assays at a time. The modifications used for the final bioassays included introduction of a loosely woven cotton cloth for application of the feeding-deterrent compounds, a collagen membrane, and use of a cocktail diet containing a red food dye (*24*) instead of blood (*30*).

**Figure 1.**
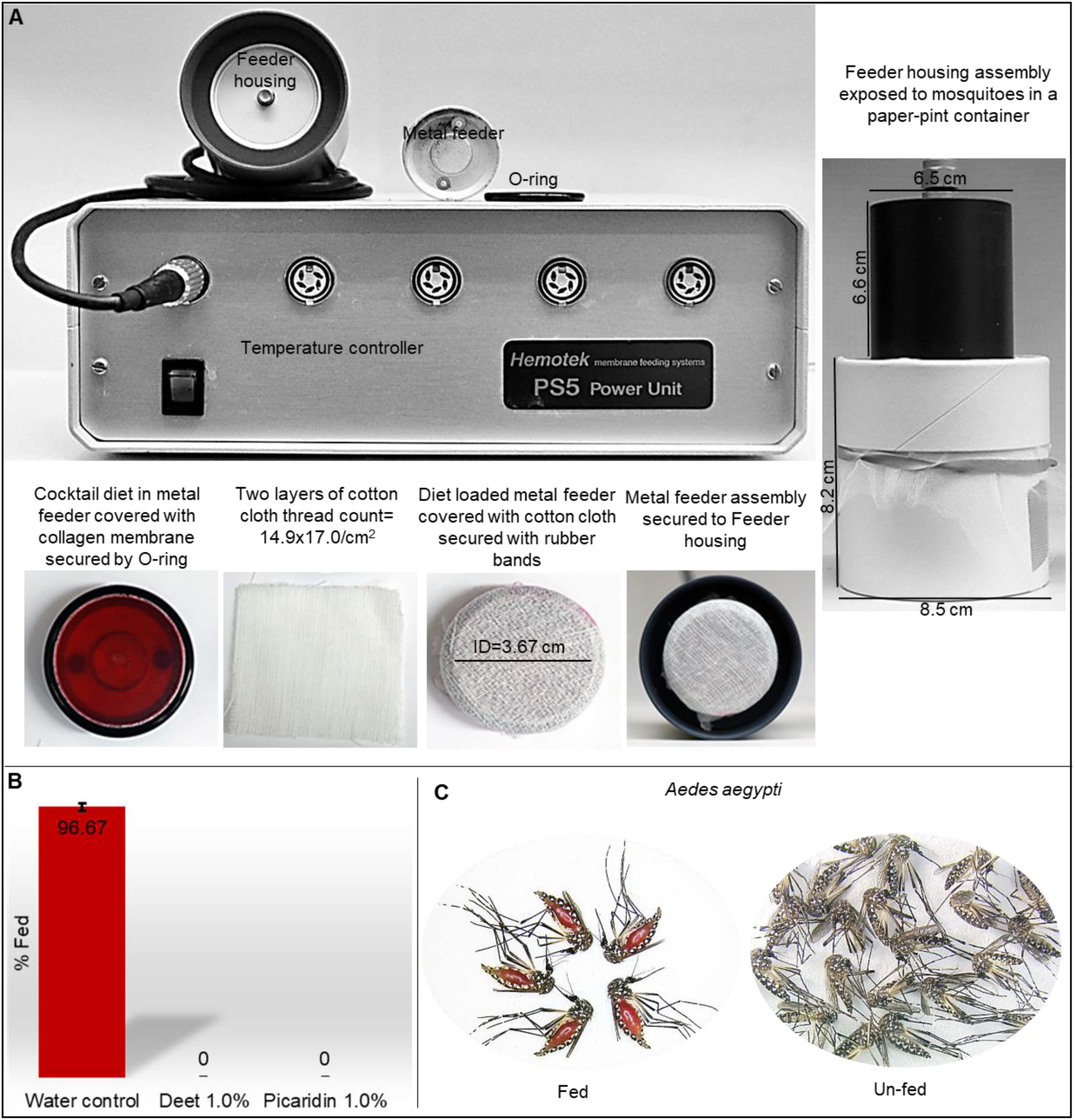
Description of the membrane feeding and feeding-deterrent screening system. Panel A shows components of the feeding system including: (Top/bottom panel L-R): Hemotek temperature controller, Feeder-housing assembly, Metal feeder assembled with cocktail diet (red color) secured with collagen casing and O-ring, Two-layers of cotton cloth, Metal feeder with cotton cloth secured via rubber bands (not visible), Metal feeder assembly secured to feeder housing. Panel B and C shows results of the feeding assays. (B) ~96% feeding rate was obtained with *Aedes aegypti* with water applied to the cotton cloth as control and 0% with 0.95 mg/cm^2^ Deet or picaridin (equivalent to 1.0% v/v) in three replicate experiments. Error bars=SEM. (C) Representative image depicting appearance of fed (red abdomens) vs un-fed *Aedes* mosquitoes resulting from the bioassay. Images clearly show engorged abdomens and red dye in fed mosquitoes. Absence of color or engorgement of abdomens indicates no feeding occurred with 1% Deet (or picaridin) as a positive control.

We first tested this system with positive (Deet or picaridin at 0.95 mg/cm^2^ Deet or picaridin in water; equivalent to 1.0% v/v) and negative (water) controls applied to cloth covering membrane feeder. Mosquitoes were allowed to feed for 30 min and then frozen at −20 °C before scoring and counting. Mosquito-feeding rate was then scored by counting fed and unfed mosquitoes following bioassays in the presence or absence of the repellent compounds. The outcomes of the screening assay are shown in Fig. 1B and C. Using this system, we obtained reproducible and consistently high mosquito-feeding rates when the cotton cloth was treated with water and 0% feeding when Deet or picaridin were tested as positive repellent controls. Results of three replicate experiments (n=20 mosquitoes per replicate; total n=60) are presented in Fig. 1B and show an average feeding rate of 96.67% (error bar=SEM) with water controls with *Aedes. Anopheles gambiae* and *Culex pipiens* also fed well (75-80%, Fig. 3) indicating that the bioassay can be used with multiple species. Based on these results, we determined that the bioassay provided a robust and reproducible test arena to screen MFD-activities in the Xbu extracts at various stages of purification.

In addition, this bioassay was optimal to screen minimal amounts of compounds, a major concern when working with naturally produced microbial compounds not easily obtainable in large quantities. The method also provided rapid results as well as consistent measures of feeding deterrence that were reproducible across assay dates.

### Purification and identification of MFD-active compounds from *X. budapestensis*

Next, we developed a procedure to isolate MFD-active compounds from Xbu cultures.

*Xenorhabdus* bacteria are known to produce antibiotics and secondary metabolites in the late stationary phase of their growth cycle (*11, 31, 32*). Accordingly, we used 72 h bacterial cultures to harvest mosquito feeding-deterrent compounds from the cell free culture supernatants. MFD-active compounds in the culture supernatants were concentrated via acetone precipitation. In the next step, water-soluble acetone precipitates yielded a broad peak detected at 280 nm on a reverse phase flash-C18 chromatography column indicating that the MFD-active compounds co-eluted with other compounds detectible at 280 nm. The MFD-activity concentrated and eluted as a single broad peak fraction. Representative image of flash-C18 purification results is shown in Supplemental Information, Fig. S3. The sample from the peak fraction was subjected to mass spectrometry analysis for determination of molecular masses and subsequent identification. MALDI-TOF revealed the presence of several abundant species including compounds at m/z 1302.94, 1312.96, and 1430.07 in this fraction (Fig. 2A) with additional masses at 44.03 m/z units higher, leading to the idea that these species were likely related. The difference of 44 arises due to addition of one C_2_H_4_O moiety (*11*).

**Figure 2.**
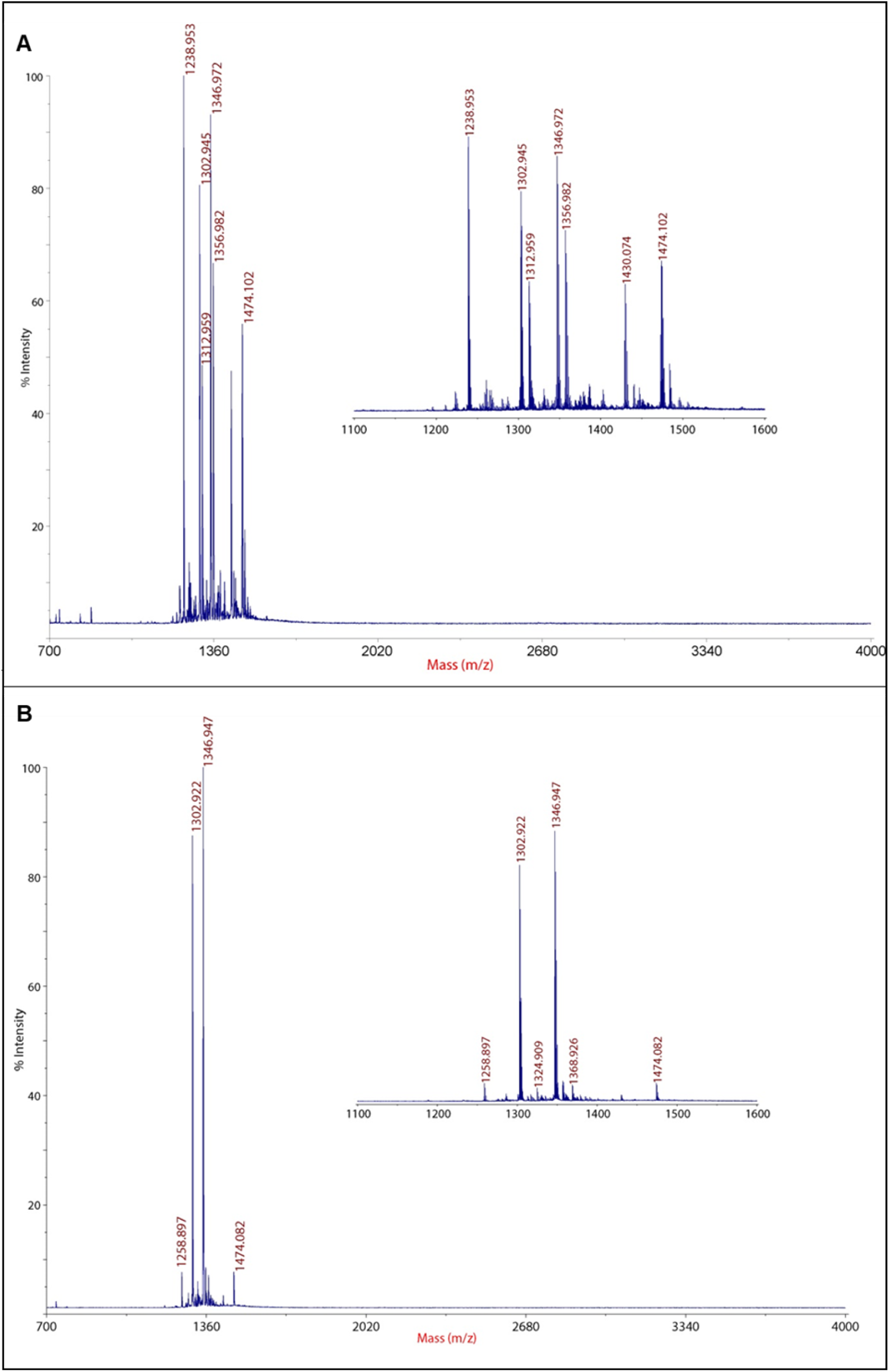
MADLI-TOF spectrum of flash & HPLC C18 reverse phase chromatography separated MFD-active peak fractions. (A) MALDI-TOF analysis of the flash-C18 purified MFD-active fraction yielded several major masses (and related species) 1302.94 (1346.97), 1312.96 (1356.98), and 1430.07 (1474.10), as well as m/z 1238.953. The mass difference of 44 m/z in these pairs has been attributed to addition or loss of C_2_H_4_O moieties (*11*). (B) MALDI-TOF analysis of HPLC-purified MFD-active, Xbu Peak#3 shows enrichment of two abundant and closely related masses at m/z 1302.92 and 1346.95. The same mass difference of 44.03 is also seen in this mass spectrum.

The MFD-active fraction from the C18 flash chromatography was passed through a 5 MWCO ultracentrifugation filter (Sartorius, Fisher Scientific, USA) and the flow through was subjected to a second purification step using HPLC on an analytical reverse phase C18 column. A gradient of 13-40% acetonitrile (ACN) over 40 min yielded four major peaks detectible at 214 nm. All of the repellent activity was concentrated in peak number 3 (hereafter referred as Xbu Peak#3), which eluted at an ACN concentration of ~24% and retention time ~14:00 min. Representative image of HPLC-C18 purification is shown in Fig. S4, with MFD-active Xbu Peak#3 indicated. Several HPLC runs were conducted to generate material for MFD-activity analysis. Subsequent MS analysis on the HPLC purified, MFD-active Xbu Peak#3, demonstrated the presence of two highly abundant masses at m/z 1302.9 and 1346.9 (Fig. 2B) indicating that these had been enriched in the MFD-active fractions as these same molecular species were observed in the first round of purification on flash C18 column. Mass spectrometry on MFD-active Peak#3 was performed on three different batches of the purified compounds. In each replicate, results were consistent, and these two masses were the highest abundance ions detected. Representative MS spectra for these experiments are presented in Fig. S5.

Since we were interested in identifying the compound/s, we subjected the MFD-active fraction to further mass spectrometry as well as total amino acid and N-terminal sequencing analysis. The active fraction was analyzed by unassisted nanospray on an Orbitrap Elite mass spectrometer (Thermo Fisher Scientific, San Jose, CA). The sample was loaded into coated, pulled borosilicate glass tips (New Objective, Woburn, MA) and after initiation of electrospray at 1.5kV, the spray was self-sustaining via capillary action. MS data was collected in profile mode in the Orbitrap analyzer at a resolving power setting of 120,000. Nanospray MS revealed the presence of the same major species observed by MALDI-MS with each compound in a dominant charge state of 2. Charge states from 1 to 4 were detected (Fig. S6). MS/MS was performed on the doubly charged species at m/z 651.9 and 673.9 using both CID and HCD fragmentation at 35% normalized collision energy. The fragmentation spectra showed that these compounds were structurally very similar, with only the difference of 44 Da between them, as previously observed by MALDI MS/MS. The MS/MS spectra and the fragment ion assignments show excellent agreement with the recently described “fabclavines” isolated from a similar strain of *Xenorhabdus budapestensis* (*11*). Using the naming convention described in that work and the structures for fabclavines Ib (m/z 1346) and IIb (m/z 1302), all major ions present in the MS/MS spectra obtained on purified Xbu compounds in this study could be accounted for (Fig. S7A and S7B).

Results of N-terminal Edman degradation analyses on MFD-active Peak#3 that contains two related peptides (provided in Fig. S8) were inconclusive in determining the sequence of amino acids, possibly explained by an inaccessible N-terminus of the molecule due to macrocylization (*11*). Total amino acids were measured from MFD-active Peak#3 by two different strong cation ion exchange chromatography methods, sodium and lithium-based elution systems (Molecular Structure Facility, University of California, Davis, CA, USA). Both systems resulted in significant signals for amino acids Asx and His (Fig. S9A and B). In addition, two unidentified signals were observed which could be 2, 3-diaminobutyric acid. Comparison with the elution profile of an available L-2, 4-diaminobutyric dihydrochloride (DAB) standard did not confirm the identity of the unknown peaks. Although the elution profiles were consistent with the expectations for DAB, DAB also co-elutes closely with other amino acids. According to the published data by Fuchs et al (*11*) fabclavines are composed of a peptide backbone containing amino acids phenylalanine or histidine, 2,3-diaminobutyric acid, three aspartic acid residues and a modified proline residue.

### MFD-activity comparison between Xbu-compounds, Deet and picaridin

Next, we determined the MFD-dose of the compounds produced by Xbu with *Aedes aegypti* mosquitoes. For this, we tested a range of concentrations of the purified MFD compounds and compared their feeding-deterrent activities to that of Deet and picaridin. Table 1A shows **f**eeding **d**eterrence **d**ose (FDD50 or FDD90) - a dose (expressed in mg/cm^2^) of the compound that resulted in 50% or 90% reduction in mosquito feeding.

FDD50 among Xbu Peak#3 fraction and Deet were similar (Xbu Peak#3 = 0.014 and Deet=0.012 mg/cm^2^) whereas it was 6.14x lower than that of picaridin (0.086 mg/cm^2^). The FDD90 value for Xbu Peak#3 was much lower compared to both Deet and picaridin (Xbu Peak#3 =0.057; Deet=0.178 and Picaridin=0.471 mg/cm^2^) indicating a much lower concentration of Xbu compounds is required to achieve FDD90 as compared to Deet or picaridin. The relative efficacy(*24*) (RE90) of Xbu peak#3 derived as a ratio of FDD90 of Deet or picaridin to that of FDD90 of Xbu Peak#3 was 3.12 and 8.26 respectively, suggesting that Xbu MFD is more potent in inhibiting mosquitoes from feeding on the cocktail diet under the conditions of the screening assay. Table 1B shows the Probit analyzed comparison among the least square estimates (*33*) for each compound’s effect. The repellency comparison based on adjusted p-values among Deet and picaridin was significant (p<0.05); Deet and Xbu Peak#3 were not significantly different (p>0.05); and Xbu Peak#3 was significantly different from picaridin (p<0.05).

**Table 1A.** Determination of feeding-deterrence dose (FDD50* and FDD90*) of Deet, picaridin and Xbu-compounds against *Aedes aegypti*.

| Compound | Slope<br>±SE | FDD50<br>(mg/cm<sup>2</sup>) | 95%<br>Fiducial<br>limits | FDD90<br>(mg/cm<sup>2</sup>) | 95%<br>Fiducial<br>limits | RE90 |
| --- | --- | --- | --- | --- | --- | --- |
| Deet | 1.09±0.36 | 0.012 | 0.000-0.032 | 0.178 | 0.066-6.785 | 1 |
| Picaridin | 1.73±0.35 | 0.086 | 0.044-0.156 | 0.471 | 0.237-2.003 | 0.38 |
| Xbu Peak#3 | 2.08±0.36 | 0.014 | 0.010-0.020 | 0.057 | 0.035-0.149 | 3.10 |

\*FDD50 and FDD90 – Concentration (expressed as mg/cm<sup>2</sup>) of the compound producing 50 or 90% reduction in feeding. RE90 - Relative efficacy derived as a ratio of FDD90 of Deet to those of picaridin and Xbu Peak#3.

**Table 1B.**
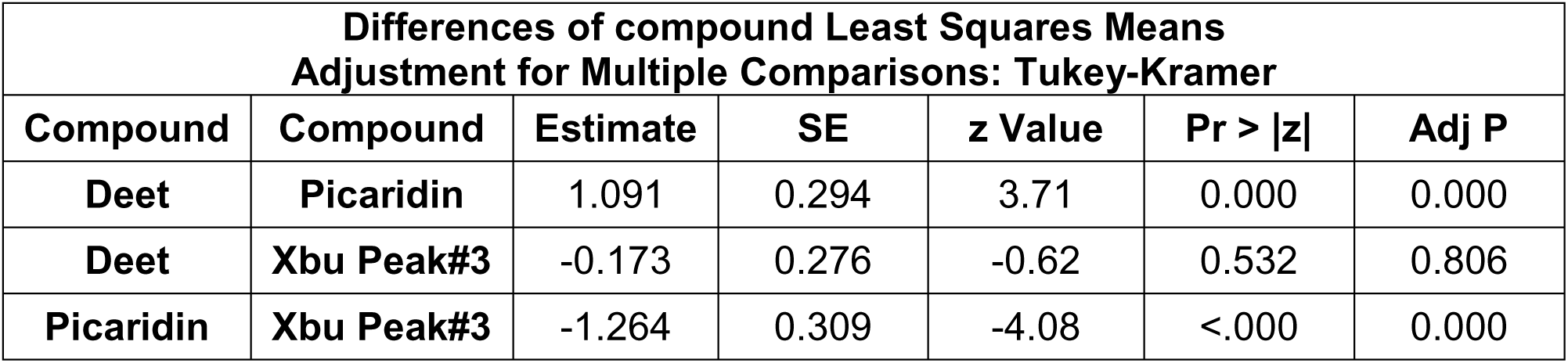
Comparison of MFD-activity of Deet, picaridin and Xbu Peak#3 against *Aedes aegypti*. Table shows the Probit analyzed comparison among the least square estimates for each compound’s effect(*33*). Feeding-deterrence comparison based on adjusted p values among Deet and picaridin was significant (Adj P<0.05); Deet and Xbu Peak#3 was not significant (Adj P>0.05); and Xbu Peak#3 was significantly different from picaridin (Adj P<0.05).

| Differences of compound Least Squares Means<br>Adjustment for Multiple Comparisons: Tukey-Kramer |  |  |  |  |  |  |
| --- | --- | --- | --- | --- | --- | --- |
| Compound | Compound | Estimate | SE | z Value | Pr > \|z\| | Adj P |
| Deet | Picaridin | 1.091 | 0.294 | 3.71 | 0.000 | 0.000 |
| Deet | Xbu Peak#3 | -0.173 | 0.276 | -0.62 | 0.532 | 0.806 |
| Picaridin | Xbu Peak#3 | -1.264 | 0.309 | -4.08 | <.000 | 0.000 |

While the exact mechanism by which Xbu compounds exert mosquito feeding-deterrence is a subject for future in-depth investigations, visual observations during feeding indicated that landing and probing by *Aedes* mosquitoes on the cloth treated with Xbu compounds was dose dependent. At a FDD50, a majority of mosquitoes landed and about half succeeded in feeding. At a FDD90 or a higher dose, few mosquitoes landed and none fed. The mosquitoes that landed attempted to probe through the cloth and were seen cleaning their proboscises soon after probing but did not imbibe food.

Thus, mosquito contact with the treated surface preceded feeding-deterrence at low dose. Deciphering feeding-deterrence by olfactory or gustatory mechanisms against low-volatility compounds, including Deet and picaridin (*34*), that require a proximate contact by mosquitoes, is difficult due overlapping sensory responses (*35*). Addition of Deet to blood (*36*) deterred mosquitoes from feeding through a direct contact with the blood meal likely via gustatory response. External gustatory sensilla on antennae of Chagas vector *Rhodnius prolixus* mediate feeding deterrence to quinine or caffeine when applied to mesh covering feeding solutions (*37*). Certain fatty acids exert feeding-deterrence in *Aedes aegypti* when applied to cloth surfaces covering artificial diet (*38*). Feeding-deterrence exhibited by Xbu compounds might elicit a combination of olfactory and gustatory responses upon contact by mosquitoes with the treated cloth surfaces in a dose dependent manner.

We also observed feeding-deterrence exhibited by Xbu compounds with *Anopheles gambiae* and *Culex pipiens*. For tests with these mosquitoes, we modified our screening assay by incorporating a single layer of a muslin cloth (rather than two layers as before). These mosquitoes had difficulty feeding with the two layers of cotton cloth described for *Aedes*. One layer of muslin yielded 75-80% feeding rate (Fig. 3). Feeding assays performed using the *Aedes* FDD90 concentration (0.057 mg/cm^2^) for *An. gambiae* and *C. pipiens* are shown in Fig. 3. Results indicate that the Xbu compounds exert a potent feeding-deterrent activity against *An. gambiae* and *C. pipiens*, as was seen with *Aedes. Aedes* mosquitoes were also included in this trial using the single layer of cloth.

**Figure 3.**
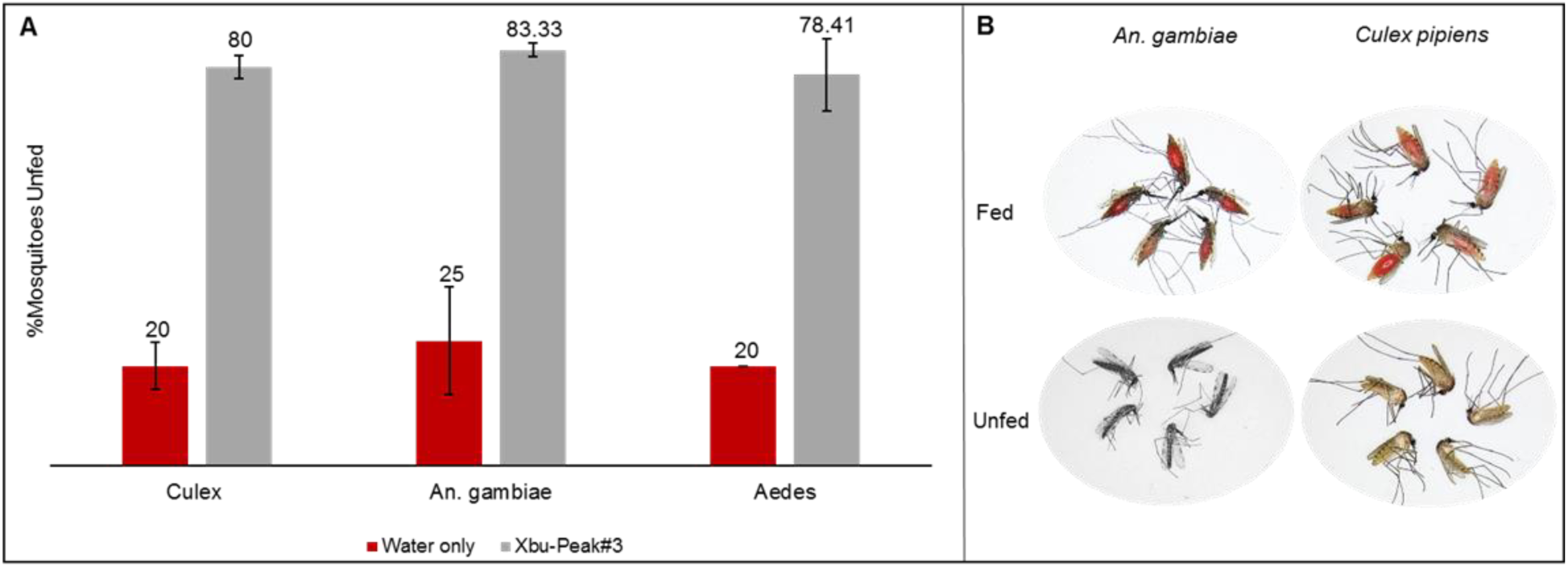
MFD-activity* of Xbu-Peak#3 with *Culex pipiens, Anopheles gambiae and Aedes aegypti*. (A) Chart shows MFD-activity of Xbu compounds tested at 0.057 mg/cm^2^. Data from three replicate experiments is shown. *Aedes aegypti* mosquitoes were included for comparison. Error bars=SEM. (B) Appearance of fed (red abdomens) and unfed Anopheles and Culex mosquitoes. *In this experiment one-layer muslin cloth was used.

## Conclusion

The bacteria associated with entomopathogenic nematodes offer an enormous potential for novel bioactivities especially attributable to low-molecular weight natural products. Genome mining projects continue to uncover information on biosynthetic gene clusters responsible for production of several of the bacterial secondary metabolites awaiting identification and assignment to their natural biological functions (*1*).

This report provides evidence that the bioactivities of the fabclavine class of natural products can be exploited in novel applications such as mosquito-feeding deterrence. Feeding deterrence activity of fabclavines determined by membrane mosquito feeding experiments is comparable to or higher than gold standard repellent Deet or picaridin against *Aedes aegypti*. Furthermore, the feeding rate of *Anopheles* and *Culex* mosquitoes decreased significantly in the presence of Xbu-compounds, suggesting that these compounds are effective in inhibiting a broader range of mosquito species. Authors recognize that the MFD-active fraction is a mixture of compounds, dominated by two co-eluting (on a reverse phase column), closely related molecular species at m/z 1302 and 1346. Future studies should 1) explore chromatographic or otherwise resolution of these compounds to identify those responsible for deterrence activity, 2) determine whether the compound(s) can affect feeding deterrence individually or require application as a mixture, 3) demonstrate deterrent activity of fabclavines via *de novo* synthesis, and 4) determine efficacy and toxicological properties of the deterrent-active compound(s) for use in mosquito deterrent/repellent formulations for eventual testing on human skin.

## Materials and methods

### Ethics statement

No live animals or human subjects were used in mosquito-deterrent screening assays in this study.

### Rearing and maintenance of mosquitoes

Colonies of *Aedes aegypti* (Liverpool strain), *Anopheles gambiae* (G3) and *Culex pipiens* (Iowa strain) were maintained at the University of Wisconsin according to standard procedures reported previously (*39-42*). Briefly, mosquito colonies were maintained at 26.5±0.5 °C and 80±5% relative humidity. Larvae were fed Tetramin^®^ fish food. Adult mosquitoes were maintained on a constant exposure to 10% sucrose presented through cotton balls. For egg production, adult female mosquitoes were offered defibrinated rabbit blood (HemoStat Laboratories, CA, USA) via Hemotek membrane feeding system (Discovery Workshops, Accrington, UK) described below. Edible collagen casing membrane (Nippi Edible Collagen Casing, ViskoTeepak, WI, USA) was selected as a membrane of choice for blood feeding and feeding-deterrent screening as all three species of mosquitoes fed readily through it. For feeding-deterrent screening assays, twenty nulliparous, mated, 7-10 day old adult female mosquitoes, hatched from a same batch of eggs were separated in screened paper containers (height x width = 8.2×8.5 cm; Neptune Paper Products, NJ, USA). Separated mosquitoes were starved for 12-16 h before the screening assays.

### Feeding-deterrent screening assay

Mosquitoes were exposed to Xbu feeding-deterrent compounds or test repellents for 30 min at room temperature (incandescent light, 25-26 °C) between 10:00 a.m. and 4:00 p.m. (USA Central time) using modified membrane-feeding set-up described in Fig. 1.

Briefly, the Hemotek temperature controller was set to a constant temperature 37 °C 5-10 min ahead of the assay. A 2.5×2.5 cm^2^ piece of a fresh, pre-cut and thoroughly water-washed collagen casing membrane was secured to the metal feeder using an O-ring. Approximately 2.5 ml of the cocktail diet^21^ containing 2% (v/v) red food dye (McCormick & Company Inc. MD, USA) was introduced from the opening at the back of the metal feeder. A cotton cloth was placed on top of the collagen casing and secured by rubber bands. Test compound or water was then applied by immersing the feeder assembly in 1 ml of the respective test solution, and the metal feeder assembly was then secured to the feeder housing and exposed to mosquitoes housed in a screened container. Several types of clothing materials varying in thickness, texture/thread count were evaluated with an objective of finding the one that provided optimal mosquito-feeding rate. Eventually, a double-layer of cotton cloth obtained from Joann Fabrics, Madison, WI, was used with *Aedes aegypti*. The features and dimensions of the chosen cotton cloth were: thread count= 14.9×17.0/cm^2^ (average of 10 measurements); diameter of the circular cloth applied to metal feeder as shown in Fig. 1A was 3.67 cm; total area of the circular cloth area exposed to mosquitoes was circa 10.57 cm^2^. However, for *Anopheles gambiae* and *Culex pipiens*, one layer of muslin cloth (thread count 24.3×33.9/cm^2^; average of 10 measurements) was used, as this cloth worked well with these mosquitoes. Both types of clothing materials are shown in Fig. S2.

Since purified Xbu compounds were dissolved in water for testing, Deet and picaridin were also prepared in 0.2 μ filtered double-distilled water. Deet was diluted from SC Johnson’s OFF! Deep Woods containing 25% Deet and picaridin from Clean Insect Repellent (Purchased from Walgreens, USA) containing 7% Picaridin. One percent stock solutions of Deet and picaridin (equivalent to 10 mg/ml) were dissolved in water from which range of dilutions was prepared for determination of feeding-deterrence dose. Both of these repellents were tested at a concentration range 1% to 0.01% (v/v) corresponding to 0.95 to 0.0095 mg/cm^2^. Both compounds dissolved well in water at the tested concentrations. Deet and picaridin were tested on the same day with four hours gap in between tests. HPLC purified, lyophilized MFD sample was dissolved in water and filtered through a 0.2 μ filter. For an accurate assessment, protein content in this sample was determined via two methods, 1) bicinchoninic acid assay according to manufacturer’s instructions (Pierce BCA Protein Assay Kit, Thermofisher Scientific, USA) and, 2) via total amino acid analysis (Molecular Structure Facility, University of California, Davis, CA). The difference in protein content determined by these two methods was negligible. Thus, for routine measurements, the BCA method was used. A dosage ranges from 0.73 to 0.048 mg/ml (v/v) corresponding to 69.5 to 4.57 μg/cm^2^ of the MFD-active compounds was tested with *Aedes aeypti*. Testing with *Anopheles gambiae* and *Culex pipiens* was done only with one concentration of Xbu compounds that yielded a feeding-deterrence dose of 90% with with *Aedes aegypti* (i.e. 0.060% v/v, corresponding to 0.057 mg/cm^2^; Table 1A). Three replicate feeding experiments were conducted for each compound with mosquitoes hatched from different egg batches (to account for cohort bias) over a period of three weeks. One replicate consisted 20 mosquitoes tested on same day with each concentration of the dilution range. For example, one replication of bacterial compound with a concentration range of 69.5 to 4.57 μg/cm^2^ was tested on same day. Extensive washing of the metal feeders, replenishing collagen membrane, cotton cloths, feeding solution for each exposure test and time gaps between assays assured that there was no carryover of the compounds/contamination of the feeders in between assays. As an extra precaution, feeding-deterrence assays with bacterial compounds were purposely performed on a different day than assays with Deet and picaridin. After the completion of the screening assay (30 min), mosquitoes were killed by freezing at −20 °C. Fed versus unfed mosquitoes were counted under a dissecting microscope and verified by at least two people. Fed mosquitoes could be easily distinguished as they were engorged and had red abdomens which were clearly visible even to an unaided eye (fully fed or partially fed) while unfed mosquitos were lean and did not have red abdomens. Mosquitoes that were not engorged and had no red color were considered as unfed and hence deterred (Fig. 1C and 3B). This distinction provided a reliable, quantitative means of assessing the feeding rate of mosquitoes with different compounds/across assays and allowed a robust comparison. Count data (proportion of unfed mosquitoes from a total) was used for statistical analysis. Several tests with HPLC purified Xbu Peak#3 from different batches exhibited consistent concentration dependent MFD-activity, in this report, results of three replications are presented. Fig. 1B, Table 1 and Fig. 3 present results of all of the feeding-deterrence screening experiments.

### Isolation, purification and identification of MFD-active compounds produced by *X. budapestensis*

*Xenorhabdus budapestensis* bacteria (*43*) obtained from Dr. Heidi-Goodrich Blair, Dept. of Bacteriology, University of Wisconsin-Madison, USA) were maintained as glycerol stocks at −80 °C. For culturing, the bacteria were freshly streaked on to NBTA plates (*43*). A single, isolated blue colony (image of bacteria streaked on LB plate is seen in Fig. S1 was taken for further growth in modified minimal medium containing 0.05 M Na_2_HPO_4_; 0.05 M KH_2_PO_4_, 0.02 M (NH_4_)_2_SO_4_, 0.001 M MgSO_4_, 0.25% yeast extract and 0.1 M glucose supplemented with 1% yeast extract. A single bacterial colony was resuspended in 1.5 ml medium from which 200 microliters were inoculated in a flask containing 400 ml medium. Bacterial cultures (typically 6 flasks of 400 ml) were grown at 30 °C at 120 rpm in a rotatory shaker to stationary phase and harvested at 72 h post inoculation via centrifugation (3913g, Beckman rotor JA14 at 4 °C).

The chilled cell free culture supernatant was mixed with two-volumes of ice-cold acetone and incubated at cold for 12-16 h while stirring. Post precipitation, spent medium/acetone was discarded and precipitates containing the MFD-active compounds were dissolved in ultrapure water so as to concentrate it to about 10x of the original culture volume (i.e. 240 ml water). This solution was first centrifuged at 3578g at 4 °C (Beckman rotor JA20) and then filtered through 0.45 μ filter and loaded on a flash C18 reverse phase column (Reveleris Flash Cartridge, 12 g, BUCHI Corporation, DE, USA) using a peristaltic pump. The column was then connected to a FPLC (Akta Prime Plus, GE Healthcare Bio-Sciences, PA, USA). Solvents for reverse phase columns were: (A) 0.2 μ filtered double distilled water, and (B) HPLC grade, 0.2 μ filtered acetonitrile (Fisher Scientific, USA). Both solvents contained 0.1% (v/v) trifluoroacetic acid (TFA).

A selected acetonitrile gradient of 50-100% was used to collect the MFD-active peak fraction on flash C18 column via FPLC. The peak fraction was lyophilized and the powered material was dissolved in ultrapure water and filtered using 0.2 μ syringe filter for downstream processing. Elution pattern of the MFD-active peak was consistent and reproducible between different batches of cultures/purifications. A representative result of this step is shown in Fig. S3.

A second reverse phase analytical column (Vydac C18, 5 μm, 4.6 mm i.d. x 150 mm); Cat# 218TP5415 currently available through Fisher Scientific, USA) was used to separate compounds enriched in the flash column purified sample. An acetonitrile gradient of 13-40% was selected. Typically, the MFD-active fractions eluted at ~ 24% acetonitrile, consistently. Multiples HPLC cycles were performed to collect enough material for downstream repellent screening and mass spectrometry analyses. A representative elution profile of MFD-active peak fraction on the HPLC column is presented in Fig. S4.

The MFD-active fractions were subjected to MALDI-MS analysis at the Mass Spectrometry Facility at the University of Wisconsin-Madison Biotechnology Center. MALDI spectra were acquired on a Sciex 4800 TOF-TOF mass spectrometer operated in positive ion reflector mode. Sample (0.5 μl) was spotted onto 384-sample Opti-TOF insert and 0.5 μl alpha-cyano-4-hydroxycinnamic acid (Fluka, Switzerland) at 6 mg/mL in 75% acetonitrile, 0.1% TFA was added and the sample and matrix were mixed on-plate. The spot was air-dried and data was acquired in positive ion reflector mode over the m/z range 700-4000 using a laser intensity of 2800 and averaging 1000 shots per spectrum. Mass assignment was performed using external calibration, with calibration taking place immediately prior to data collection. Electrospray MS and MS/MS were performed using unassisted nanospray by loading the MFD-active fraction into pulled, coated borosilicate tips and acquiring positive ion, profile mode spectra on an Orbitrap Elite mass spectrometer (Thermo Fisher Scientific, CA, USA). MS and MS/MS data were collected in the Orbitrap analyzer at a resolving power setting of 120,000. MS data was acquired over the m/z range 150-1800, while HCD MS/MS data was collected from m/z 100-1500 and CID MS/MS spectra were collected from m/z 180-1500. CID and HCD fragmentation used 35% normalized collision energy.

### Statistical analysis

Probit analysis (Statistical Analysis Software, version 9.4; SAS Institute Inc. NC, USA) on the log (base 10) transformed feeding data was used to estimate feeding-deterrence dose (FDD). Feeding-deterrent activity (Table 1) was expressed as a mg/cm^2^ dose of the compound (bacterial or Deet or picaridin) applied to cloth that resulted in a 50% or 90% inhibition in mosquito-feeding rate (FDD50 and FDD90) similarly as described previously (*24*). Relative efficacy (RE90) of the deterrent activity was derived by dividing FDD90 value of the Deet to that of the picaridin and Xbu Peak#3, respectively. Least square estimation (*33*) was conducted to compare differences among feeding-deterrence activity between three compounds based on adjusted p-values (Table 2).

## Acknowledgements

Support from NIH R21 grant # AI123719 to SMP is greatly acknowledged. We thank Dr. Heidi Goodrich-Blair and Dr. Angel Torres for sharing the strain of *Xenorhabdus budapestensis*, Dr. Il-Hwan Kim for discussions on bacterial cultures, Dr. Lyric C. Bartholomay for providing colonies of mosquitoes, Dr. Bartholomay and Dr. Robby Weyker for suggestions on collagen casing, Edmund Norris for suggestions on repellent assays, Undergraduates Nathan Wong, Sienna Muehlfed and Mara Dhyr for help with mosquito colony maintenance, Dr. Walter Goodman and Dr. Ronney Frederick for consultation on MFD-active compound purification, Yongsu Lee & Dr. Jun Zhu for support with statistical analysis.

## Author contributions

S.M.P. conceived the study, provided guidance on study design and edited manuscript. M.K.K. designed and performed experiments, collected and analyzed data and wrote the manuscript. G.A.B-W. collected all mass spectrometry data and annotated the MS/MS spectra. All authors edited, reviewed and approved the manuscript.

## Additional information

### Supplementary information

Provided as a word document.

### Data availability

The data presented in this report are available from the first or the corresponding author on request.

## References

1. K. A. J. Bozhüyük et al., Natural Products from Photorhabdus and Other Entomopathogenic Bacteria. Curr Top Microbiol Immunol 402, 55–79 (2017).

2. J. B. Gonzalez, F. J. Fernandez, A. Tomasini, Microbial Secondary Metabolites Production and Strain Improvement. Indian J of Biotechnology 2, 322–333 (2003).

3. H. B. Bode, Entomopathogenic bacteria as a source of secondary metabolites. Current opinion in chemical biology 13, 224–230 (2009).

4. R. Akhurst, Antibiotic activity of Xenorhabdus spp., bacteria symbiotically associated with insect pathogenic nematodes of the families Heterorhabditidae and Steinernematidae. Microbiology 128, 3061–3065 (1982).

5. G. Chen, G. Dunphy, J. Webster, Antifungal activity of two Xenorhabdus species and Photorhabdus luminescens, bacteria associated with the nematodes Steinernema species and Heterorhabditis megidis. Biological Control 4, 157–162 (1994).

6. X. Cai et al., Entomopathogenic bacteria use multiple mechanisms for bioactive peptide library design. Nature Chemistry 9, 379–386 (2017).

7. J. M. Crawford, C. Portmann, X. Zhang, M. B. Roeffaers, J. Clardy, Small molecule perimeter defense in entomopathogenic bacteria. Proceedings of the National Academy of Sciences 109, 10821–10826 (2012).

8. D. Reimer et al., Rhabdopeptides as Insect-Specific Virulence Factors from Entomopathogenic Bacteria. Chembiochem 14, 1991–1997 (2013).

9. I.-H. Kim et al., Specificity and putative mode of action of a mosquito larvicidal toxin from the bacterium Xenorhabdus innexi. Journal of invertebrate pathology 149, 21–28 (2017).

10. N. J. Tobias et al., Natural product diversity associated with the nematode symbionts Photorhabdus and Xenorhabdus. Nature Microbiology, 1 (2017).

11. S. W. Fuchs, F. Grundmann, M. Kurz, M. Kaiser, H. B. Bode. Fabclavines: Bioactive Peptide–Polyketide-Polyamino Hybrids from Xenorhabdus. ChemBioChem 15, 512–516 (2014).

12. M. Sergeant et al., Identification, typing, and insecticidal activity of Xenorhabdus isolates from entomopathogenic nematodes in United Kingdom soil and characterization of the xpt toxin loci. Applied and environmental microbiology 72, 5895–5907 (2006).

13. X. Zhou, H. K. Kaya, K. Heungens, H. Goodrich-Blair, Response of ants to a deterrent factor (s) produced by the symbiotic bacteria of entomopathogenic nematodes. Applied and Environmental Microbiology 68, 6202–6209 (2002).

14. B. Gulcu, S. Hazir, H. K. Kaya, Scavenger deterrent factor (SDF) from symbiotic bacteria of entomopathogenic nematodes. Journal of invertebrate pathology 110, 326–333 (2012).

15. M. S. Fradin, Mosquitoes and mosquito repellents: a clinician’s guide. Annals of internal medicine 128, 931–940 (1998).

16. F. O. Okumu, S. J. Moore, Combining indoor residual spraying and insecticide-treated nets for malaria control in Africa: a review of possible outcomes and an outline of suggestions for the future. Malaria journal 10, 208 (2011).

17. W. S. Leal, The enigmatic reception of DEET—the gold standard of insect repellents. Current opinion in insect science 6, 93–98 (2014).

18. M. Klier, F. Kuhlow, Neue Insektenabwehrmittel—am Stickstoff disubstituierte beta-Alaninderivate. J Soc Cosmet Chem 27, 141–153 (1976).

19. J. Boeckh et al., Acylated 1, 3-Aminopropanols as Repellents against Bloodsucking Arthropods. Pest Management Science 48, 359–373 (1996).

20. V. Dethier, B. L. Browne, C. N. Smith, The designation of chemicals in terms of the responses they elicit from insects. Journal of Economic Entomology 53, 134–136 (1960).

21. G. Who, Report of the WHO Informal Consultation on the evaluation and testing of insecticides. World Health Organization Geneva, (1996).

22. D. R. Barnard, Biological assay methods for mosquito repellents. Journal of the American Mosquito Control Association 21, 12–16 (2005).

23. D. Carlson, C. Schreck, R. Brenner, Carbon dioxide released from human skin: effect of temperature and insect repellents. Journal of medical entomology 29, 165–170 (1992).

24. T. H. Huang, N. Y. Tien, Y. P. Luo, An in vitro bioassay for the quantitative evaluation of mosquito repellents against Stegomyia aegypti (= Aedes aegypti) mosquitoes using a novel cocktail meal. Medical and veterinary entomology 29, 238–244 (2015).

25. J. A. Klun, M. Kramer, M. Debboun, A new in vitro bioassay system for discovery of novel human-use mosquito repellents. Journal of the American Mosquito Control Association 21, 64–70 (2005).

26. J. A. Klun, M. Kramer, A. Zhang, S. Wang, M. Debboun, A quantitative in vitro assay for chemical mosquito-deterrent activity without human blood cells. Journal of the American Mosquito Control Association 24, 508–512 (2008).

27. A. Ali, C. L. Cantrell, I. A. Khan, A New In Vitro Bioassay System for the Discovery and Quantitative Evaluation of Mosquito Repellents. J Med Entomol 54, 1328–1336 (2017).

28. T. Kröber, S. Kessler, J. Frei, M. Bourquin, P. M. Guerin, An in vitro assay for testing mosquito repellents employing a warm body and carbon dioxide as a behavioral activator. Journal of the American Mosquito Control Association 26, 381–386 (2010).

29. H. Qiu, H. W. Jun, J. W. McCall, Pharmacokinetics, formulation, and safety of insect repellent N, N-diethyl-3-methylbenzamide (deet): a review. Journal of the American Mosquito Control Association 14, 12–27 (1998).

30. A. Ali, C. L. Cantrell, I. A. Khan, A New In Vitro Bioassay System for the Discovery and Quantitative Evaluation of Mosquito Repellents. Journal of Medical Entomology, (2017).

31. G. Furgani et al., Xenorhabdus antibiotics: a comparative analysis and potential utility for controlling mastitis caused by bacteria. Journal of applied microbiology 104, 745–758 (2008).

32. J. Masschelein et al., The zeamine antibiotics affect the integrity of bacterial membranes. Applied and environmental microbiology 81, 1139–1146 (2015).

33. R. C. Littell, W. W. Stroup, R. J. Freund, SAS for linear models. (SAS institute, 2002).

34. E. J. Norris, J. R. Coats, Current and Future Repellent Technologies: The Potential of Spatial Repellents and Their Place in Mosquito-Borne Disease Control. Int J Environ Res Public Health 14, (2017).

35. J. T. Sparks, J. C. Dickens, Mini review: Gustatory reception of chemicals affecting host feeding in aedine mosquitoes. Pestic Biochem Physiol 142, 15–20 (2017).

36. W. Lu, J. K. Hwang, F. Zeng, W. S. Leal, DEET as a feeding deterrent. PLoS One 12, e0189243 (2017).

37. G. Pontes, S. Minoli, I. O. Insaurralde, M. G. de Brito Sanchez, R. B. Barrozo, Bitter stimuli modulate the feeding decision of a blood-sucking insect via two sensory inputs. J Exp Biol 217, 3708–3717 (2014).

38. A. Ali et al., Aedes aegypti (Diptera: Culicidae) biting deterrence: structure-activity relationship of saturated and unsaturated fatty acids. J Med Entomol 49, 1370–1378 (2012).

39. S. Paskewitz, S. Reese-Stardy, M. Gorman, An easter-like serine protease from Anopheles gambiae exhibits changes in transcript abundance following immune challenge. Insect molecular biology 8, 329–337 (1999).

40. M. K. Kajla, O. Andreeva, T. M. Gilbreath, S. M. Paskewitz, Characterization of expression, activity and role in antibacterial immunity of Anopheles gambiae lysozyme c-1. Comparative Biochemistry and Physiology Part B: Biochemistry and Molecular Biology 155, 201–209 (2010).

41. B. M. Christensen, D. R. Sutherland, L. N. Gleason, Defense reactions of mosquitoes to filarial worms: comparative studies on the response of three different mosquitoes to inoculated Brugia pahangi and Dirofilaria immitis microfilariae. Journal of invertebrate pathology 44, 267–274 (1984).

42. M. T. Aliota, S. A. Peinado, J. E. Osorio, L. C. Bartholomay, Culex pipiens and Aedes triseriatus mosquito susceptibility to Zika virus. Emerging infectious diseases 22, 1857 (2016).

43. K. Lengyel et al., Description of four novel species of Xenorhabdus, family Enterobacteriaceae: Xenorhabdus budapestensis sp. nov., Xenorhabdus ehlersii sp. nov., Xenorhabdus innexi sp. nov., and Xenorhabdus szentirmaii sp. nov. Systematic and applied microbiology 28, 115–122 (2005).

